# MEGSA: A powerful and flexible framework for analyzing mutual exclusivity of tumor mutations

**DOI:** 10.1101/017731

**Authors:** Xing Hua, Paula L. Hyland, Jing Huang, Bin Zhu, Neil E. Caporaso, Maria Teresa Landi, Nilanjan Chatterjee, Jianxin Shi

## Abstract

The central challenge in tumor sequencing studies is to identify driver genes and pathways, investigate their functional relationships and nominate drug targets. The efficiency of these analyses, particularly for infrequently mutated genes, is compromised when patients carry different combinations of driver mutations. Mutual exclusivity analysis helps address these challenges. To identify mutually exclusive gene sets (MEGS), we developed a powerful and flexible analytic framework based on a likelihood ratio test and a model selection procedure. Extensive simulations demonstrated that our method outperformed existing methods for both statistical power and the capability of identifying the exact MEGS, particularly for highly imbalanced MEGS. Our method can be used for *de novo* discovery, pathway-guided searches or for expanding established small MEGS. We applied our method to the whole exome sequencing data for fourteen cancer types from The Cancer Genome Atlas (TCGA). We identified multiple previously unreported non-pairwise MEGS in multiple cancer types. For acute myeloid leukemia, we identified a novel MEGS with five genes (*FLT3, IDH2, NRAS, KIT* and *TP53*) and a MEGS (*NPM1, TP53* and *RUX1*)s whose mutation status was strongly associated with survival (*P*=6.7×10^-4^). For breast cancer, we identified a significant MEGS consisting of *TP53* and four infrequently mutated genes (*ARID1A, AKT1, MED23* and *TBL1XR1*), providing support for their role as cancer drivers.

## Background

Cancers, driven by somatic mutations, cause over eight million deaths worldwide each year. Recent technical advances in next generation sequencing and bioinformatic analyses have greatly advanced the characterization of tumor genomes. Large-scale cancer genomics projects, e.g. the Therapeutically Applicable Research to Generate Effective Treatments (TARGET) for childhood cancers, The Cancer Genome Atlas (TCGA) and the International Cancer Genome Consortium (ICGC) for adult cancers, have accumulated a large amount of multi-dimensional genomic data for dozens of cancers. The primary aim in analyzing these unprecedented “big” genomic data is to identify “driver” mutation events related with tumor initiation and progression. Typically, driver genes are nominated by examining whether the non-synonymous mutation rate exceeds the background silent mutation rate^1,2^. However, identifying infrequently mutated driver genes requires a very large sample size to achieve statistical significance. A closely related challenge is to investigate relationships among mutated genes and to identify oncogenic pathways. Mutual exclusivity (ME) analysis is an effective computational approach that helps address both problems.

ME analysis was initially proposed for pairs of genes and has produced important findings that have been consistently replicated, e.g. ME between *EGFR* and *KRAS* in lung adenocarcinoma^3-5^. Because cancer pathways typically involve multiple genes, recent methods^6-11^have tried to extend pair wise analyses to search for mutually exclusive gene sets (MEGS), which also has much better power than pair wise analyses. Briefly, given a somatic mutation matrix for *N* patients and *M* genes, we aim to identify “optimal” gene subsets that are mutually exclusive.

Multiple methods have been proposed for ME analysis. Dendrix^7^ and two other methods^9,10^ use a “weight” statistic as the criterion to search for MEGS. However, this statistic is inappropriate to compare gene sets and tends to identify large MEGS with many false positive genes, as we will show in simulations. MEMo^6^ uses external biological data to form “cliques” (fully connected gene networks) and searches for MEGS within each clique to increase power by reducing multiple testing. As we will demonstrate, MEMo results in incorrect false positive rates for each clique and tends to select MEGS with false positive genes. Szczurek and Beerenwinkel^8^ proposed a non-standard likelihood ratio test but ended up with a severely misspecified null distribution. Mutex^11^ has improved existing methods and used permutations to control false positive rates; however, its overly simple statistic warrants further improvement. In summary, most of the existing methods fail to correctly control for false positive rates and lack a criterion for selecting “optimal” MEGS. Since some of these MEGS methods have been widely used in tumor sequencing projects, previous results may need to be interpreted with caution.

Ideally, an analytic framework for identifying MEGS shall have the following components. First, given a subset of *m* (*m* ≤*M*) genes, a statistically powerful test is required to examine whether mutations in these *m* genes show ME. Second, it is crucial to determine whether any subset of the *M* genes is statistically significant after adjusting for multiple testing. Third, a model selection criterion is required to compare nested gene sets to select the “optimal” MEGS. An inappropriate criterion may falsely include genes into MEGS or exclude true genes from MEGS.

We developed a framework that fits all above requirements. We developed a likelihood ratio test (LRT) for testing ME and performed a multiple-path linear search together with permutations to test the global null hypothesis, *i.e.* the set of *M* genes does not contain MEGS of any size. When global null hypothesis was rejected, we proposed a model selection procedure based on permutations to identify “optimal” MEGS. All algorithms have been implemented in an R package called MEGSA (<u>M</u>utually <u>E</u>xclusive <u>G</u>ene <u>S</u>et <u>A</u>nalysis). Extensive simulations demonstrated that MEGSA outperformed existing methods for *de novo* discovery and dramatically improved the accuracy of recovering exact MEGS, particularly for imbalanced MEGS. MEGSA can either be used for *de novo* discovery or by incorporating existing biological datasets (e.g. KEGG pathways and protein-protein interactions) to improve statistical power by reducing multiple testing, in spirit similar to MEMo^6^ and Mutex^11^. We can also use MEGSA to expand well-established small MEGS with further improved power.

We applied MEGSA to analyze the whole exome sequencing data of 14 cancer types from TCGA. We identified multiple significant non-pairwise MEGS for breast cancer, low grade glioma, uterine corpus endometrial carcinoma skin cutaneous melanoma, head and neck squamous cell carcinoma and acute myeloid leukemia with important biological implications. Incorporating KEGG pathway data further identified 8 MEGS for breast cancer and 10 for low grade glioma. Although *de novo* discovery has lower power due to the high multiple testing burden, it has the potential to identify a more complete MEGS. Incorporating external information may identify significant but likely incomplete oncogenic pathways. Thus, MEGSA shall be applied using these complimentary search strategies. We expect MEGSA to be useful for identifying oncogenic pathways and driver genes that would have been missed by frequency-based methods. MEGSA is freely available at http://dceg.cancer.gov/tools/analysis/MEGSA.

## 2. Results

### 2.1 MEGSA: a framework for identifying mutually exclusive gene sets

We consider a binary mutation matrix *A* with *N* rows (cancer patients) and *M* columns (genes), where each row represents the mutational status for one patient and each column for one gene (Fig. 1A). Let *a*_*ik*_ denote the mutation status with *a*_*ik*_=1 if gene *k* is somatically mutated for patient *i* and *a*_*ik*_=0 otherwise. Here, a somatic mutation could be copy number alternations, non-synonymous point mutations or point mutations predicted to be deleterious. We consider non-synonymous point mutations in the manuscript. MEGSA has three components: (1) an efficient likelihood ratio test (LRT) for examining ME for a subset of genes; (2) a multiple-path linear search algorithm and a permutation framework to evaluate the global null hypothesis (GNH) and (3) a model selection procedure to identify the “optimal” MEGS.

**Figure 1.**
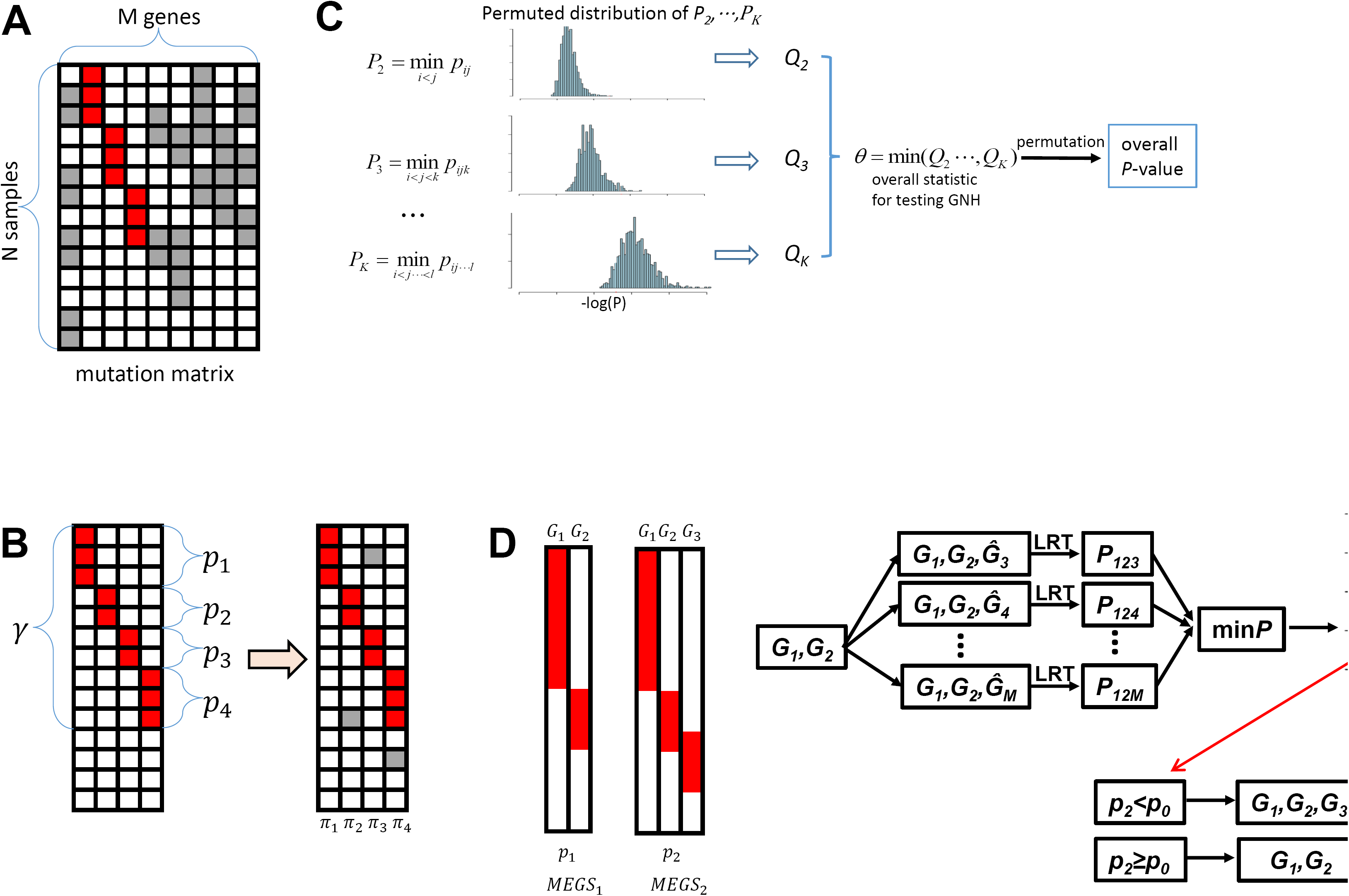
Overview of the algorithms implemented in MEGSA for searching mutually exclusive gene sets. (A) Observed somatic mutation matrix. Each row is for one sample and each column for one gene. Red entries represent MEGS mutations while gray entries represent background mutations. (B) A data generative model for MEGS. The left panel shows an MEGS with four genes showing complete mutual exclusivity. The right panel shows MEGS mutations and background mutations. γ is the coverage of the MEGS, defined as the proportion of samples covered by the MEGS. (*p*_*1*_,…,*p*_*m*_) are the relative mutation frequencies normalized to have *p*_*1*_ +… + *p*_*m*_ = 1. (C) Overall statistic for testing global null hypothesis and its significance. *p_ij_* is the *P*-value of our LRT for a gene pair (*i, j*). *p*_*ijk*_ is the *P*-value a gene triplet (*i, j, k*). For each *k*, let *P*_*t*_ as the minimum *P*-value of all sets of *k* genes and evaluate its significance (denoted as *Q*_*t*_) using permutations preserving mutational frequencies. The overall statistic is defined as *θ*=min (*Q*_2_…,*Q*_*k*_) and its significance is assessed by permutations. (D) Model selection based on permutations. Two nested MEGS (*G_1_,G_2_*) and (*G_1_,G_2_,G_3_*) have nominal P-value *pi* and *p*_2_ based on LRT. We permute mutations in (*G_3_,…,G_M_*) *by* keeping the mutual exclusivity of (*G*_*1*_, *G*_*2*_) unchanged. *G*_*t*_ represents permuted mutations. For each permutation, we calculate the minimum *P*-value for all *M-2* triplets (*G*_*1*_,*G*_*2*_,*G*_*k*_). Threshold *p*_0_ is chosen at level 5%.

#### 2.1.1 A likelihood ratio statistic for testing mutual exclusivity

Given a subset of *m (m* ≤ *M*) genes and the binary mutation matrix (denoted as *A*_0_, a sub matrix of *A*), we describe a data generative model for MEGS. We assume that the *m* genes in the MEGS are completely mutually exclusive with coverage denoted as γ, defined as the proportion of samples covered by the MEGS.Within the MEGS, we assume (*p*_*1*_,…, *p*_*m*_) as the relative mutation frequencies with *p*_*1*_ +…+ *p*_*m*_ = 1. We assume that the observed mutation matrix *A*_*0*_ is generated in three steps (Fig. 1B):

(1) Given *N* patients and coverage γ, we randomly sample *n* patients coved by the MEGS according to the distribution *Bionomial(N, γ)*.
(2) For each sampled patient covered by the MEGS, we randomly choose a “mutated” gene according to(*p*_*1*_,…, *p*_*m*_)
(3) Independent of the MEGS, we randomly simulate background mutations to each entry of matrix *A*_*0*_ with gene-specific background mutation rates ∏ = (*π*_1_…*π*_*m*_).

Based on this data generative model and further assuming *p*_*k*_ ∝ *π*_*k*_, the log likelihood is given as

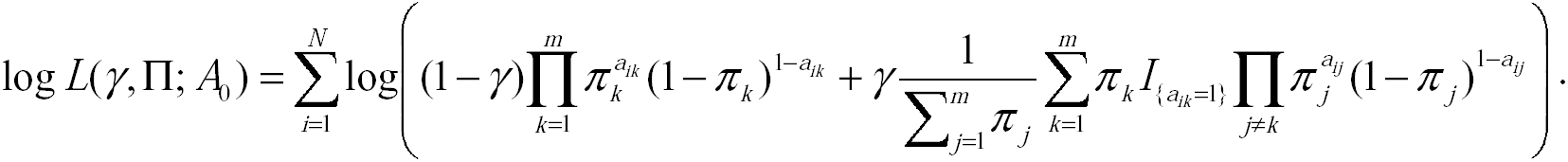

Here, γ=0 corresponds to the null hypothesis that the *m* genes are randomly mutated. Π = (*π*_1_,…,*π*_*m*_) are nuisance parameters. LRT can be derived to test *H*_*0*_: γ= 0 *v.s. H*_*1*_: γ *>* 0. Asymptotically, LRT has a null distribution 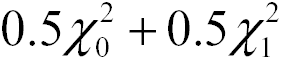,a mixture distribution with 0.5 probability at point mass zero and 0.5 probability as 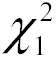 See **Methods** for details.

#### 2.1.2 Testing the global null hypothesis

Given a mutation matrix *A* with all *M* genes, it is crucial to test GNH that all genes are mutated independently. Suppose that we are interested in MEGS with no more than *K* genes. We have 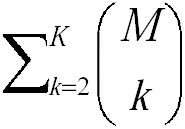combinations of genes to be tested, which equals to 2.0×10^11^ if *M*=100 and *K*=8. The multiple testing burden increases with size exponentially when *K*<*M/2.* Importantly, the total multiple testing burden is dominated by the largest MEGS with *K* genes. When *M*=100 and *K*=8, the number of tests for MEGS with 8 genes account for 91.5% of total 2.0×10^11^ tests while such proportion is only 8.0×10^-5^% for MEGS of 3 genes. Intuitively, for the same nominal *P*-value of 10^−6^, a MEGS with 3 genes should be much more significant than the one with 8 genes. Thus, putative MEGS of different sizes must be differentially treated. Moreover, the statistic tests may be highly correlated; thus the Bonferroni correction is too conservative. We propose a permutation-based procedure to address these problems **(Fig. 1C).** Note that permutations were performed by preserving the mutation frequency for each gene^7,11^.

Briefly, we first perform multiple test correction separately for MEGS of each size. For a given *k (k ≤ K),* we search all gene sets of size *k* to test for ME using our LRT and denote the minimum *P*-value as *P*_*k*_. The significance of *P*_*k*_, (denoted as Q_k_) is estimated by permutations. Because we search for MEGS of different sizes, the overall statistic for testing GNH is θ=min (*Q_2_,…, Q_K_*), with significance evaluated by permutations. Finding the minimum *P*-value *P*_*k*_ by exhaustive search is computationally challenging even for a moderate *k*. Thus, we implemented a multiple-path linear search algorithm to approximate *P*_*k*_(**Methods**).

#### 2.1.3 Identifying optimal mutually exclusive gene sets using model selection

When GNH is rejected, we can use the multiple-path linear search algorithm to identify all significant putative MEGS (**Methods).** These putative MEGS may be nested. Consider two significant putative MEGS: MEGS1 has two genes (*G*_1_, *G*_2_) with nominal *P*-value *p*_*1*_ and MEGS2 has three genes (*G*_1_, *G*_2_, *G*_3_) with nominal *P*-value *p*_2_ based on LRT. Intuitively, if *p*_*2*_≪*p*_*1*_, we choose MEGS2 with three genes. However, a simple criterion *p*_*2*_ <*p*_*1*_ is too liberal and tend to include *G*_*3*_ into MEGS even if *G*_*3*_ is independent of (*G*_*1*_,*G*_*2*_). This is because *G*_*3*_ is chosen from the *M-2* genes (*G*_3_,…,*G*_*M*_) to form the strongest MEGS with *G*_*1*_ and *G*_*2*_.

We addressed the problem in a statistical testing framework (Fig. 1D). The null hypothesis is that none of the *M-2* genes (*G*_3_,…,*G*_*M*_) is mutually exclusive of (*G*_1_, *G*_2_). We reject the null hypothesis (and thus choose MEGS2) if *p*_2_<*p*_0_ with *p*_0_ chosen to control false positive rate <5% based on permutations. Note that we keep the relationship between *G*_*1*_ and *G*_2_ unchanged and permute mutations only in (*G*_3_,…,*G*_*M*_). If (*G*_3_,…,*G*_*M*_) are independent of (*G*_1_,*G*_2_), using *p*_0_ as threshold will correctly choose MEGS1 with probability 95%.

#### 2.1.4 Identifying mutually exclusive gene sets using three search strategies

We propose three complimentary strategies for searching MEGS using MEGSA, as illustrated in Fig. 2. The first strategy is *de novo* discovery by directly applying MEGSA to all *M*genes (Fig. 2A). The advantage of *de novo* discovery is that it does not rely on any prior information and has the potential to identify a complete MEGS. However, *de novo* analyses may have low power because of heavy multiple testing burden.

**Figure 2.**
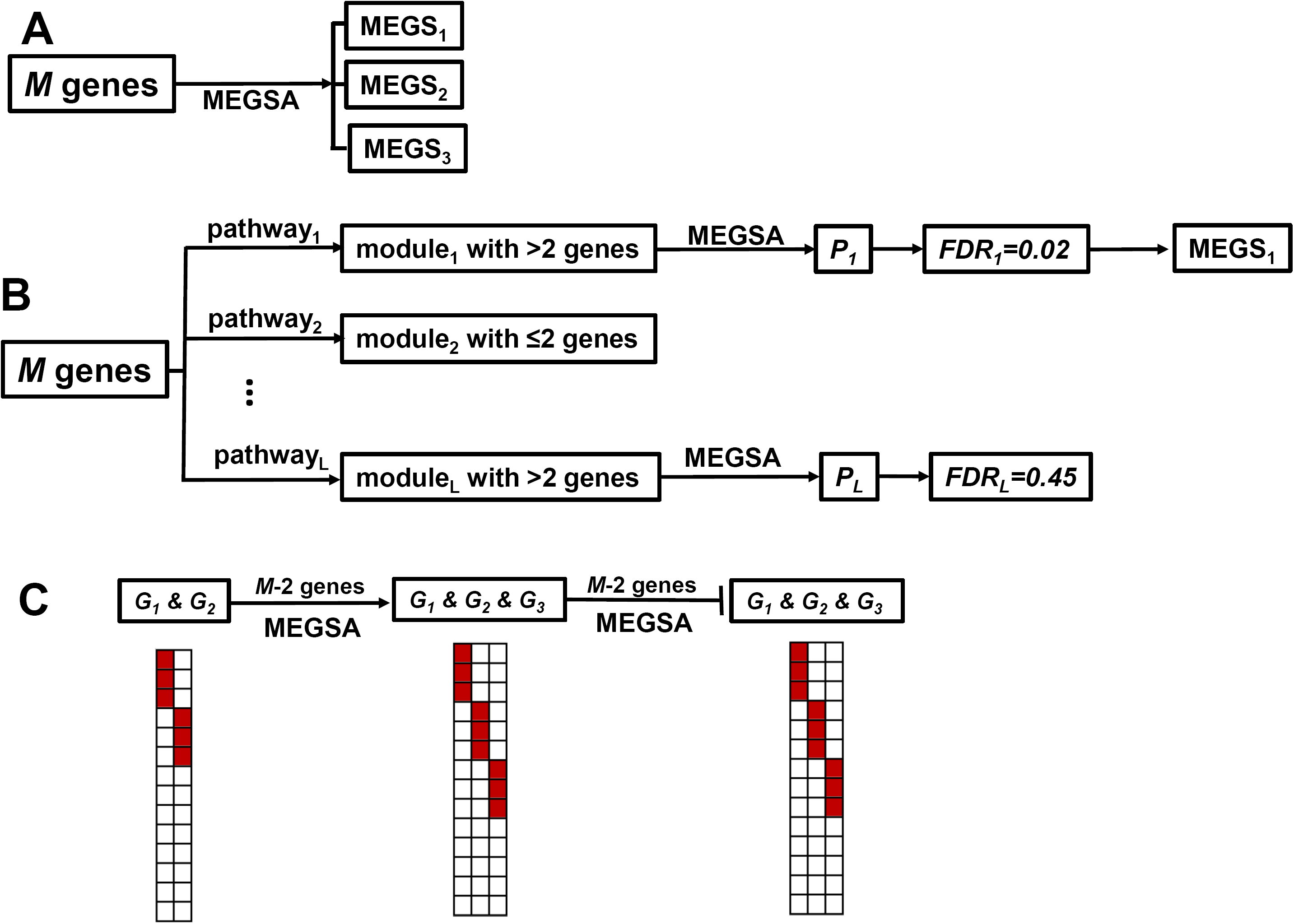
Three strategies for searching mutually exclusive gene sets using MEGSA. (A) *de novo* analyses for all *M* genes. (B) Search MEGS by incorporating KEGG pathways. For each pathway, we derive the subset (called a module) of *M* genes (with mutation data) in the pathway. We eliminate duplicate modules or modules with less than 3 genes, and analyze each module using MEGSA to derive module-wise *P*-values. We control FDR<0.05 using these module-wise *P*-values to choose significant modules and identify optimal MEGS. (C) Expanding established small MEGS (*G*_1_, *G2*) by the model selection procedure described in Fig. 1D.

MEGSA can also be applied by incorporating existing biological data, in spirit similar to MEMo^6^ and Mutex^11^. MEMo searches for fully-connected sub graphs (called “cliques”) using existing pathway and functional information (e.g. protein-protein interaction and gene coexpressoin) and analyzes each clique. Mutex restricts search space so that genes in MEGS have a common downstream signaling target. Although MEGSA can be modified to perform similar search, we exemplify this approach by using the KEGG pathway database (Fig. 2B). Briefly, we compare *M* genes with KEGG pathways and identify subsets (called modules) with more than 2 genes. We analyze each module using MEGSA and produce an overall *P*-value. We choose significant modules by controlling FDR at 5%.

The third strategy is to search MEGS starting with a well-established small MEGS (e.g. *EGFR* and *KRAS* in lung cancer). We use our model selection procedure (Fig. 1D) to “grow” the MEGS until no gene can be included (Fig. 2C).

### 2.2 Evaluation on simulated cancer mutation data

#### 2.2.1 Type-I error rate and power behavior of LRT

Since the LRT is the foundation for our algorithm, we first evaluated its type-I error rate and the power behavior for a fixed set of *m* genes. Under *H*_0_, LRT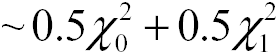 asymptotically. Results based on 100,000 simulations verified that the *P*-values calculated based on the asymptotic distribution agreed well with the simulation-based *P*-values (**Supplemental Table 1**) for different combinations of parameters, including background mutation rate, sample size and the size of gene sets. The power of LRT increases with sample size and coverage and reduces with background mutation rates (**Supplemental Fig. S1**).

#### 2.2.2 Comparison with other methods that detect mutually exclusive gene mutations

We compared the performance of MEGSA with the performances of existing methods including RME^12^, MEMo^6^, Dendrix^7^, LRT-SB^8^ and Mutex^11^. MDPFinder^9^ uses the same “weight” statistic as Dendrix but a more efficient computational method for searching MEGS; thus the comparative study does not include MDPFinder. A systematic comparison is very difficult for following reasons. Dendrix, RME and LRT-SB perform *de novo* analyses; MEMo uses existing biological data to reduce the search space; while MEGSA and Mutex can perform both analyses. In addition, for RME, Dendrix and LRT-SB, it is unclear how multiple testing was corrected. Mutex^11^ compared the performances using receiver operating characteristic (ROC) analysis; however, it is unclear how false positives and false negatives were calculated. A more detailed summary and critique of these methods can be found in the **Supplemental Note.**

We empirically evaluated the null distribution of LRT-SB^8^. Simulation results show that the empirical distribution of LRT-SB deviates dramatically from the claimed null distribution N(0,1)(**Supplemental Fig. S2**). See also the theoretical explanation in **Supplemental note**. MEMo derives a *P*-value for each “clique” and selects significant cliques by controlling FDR using these *P*-values. Controlling FDR requires *P*-values for null statistics to follow a uniform distribution U[0,1]^13^. However, our simulation results (under H_0_) show that the *P*-values dramatically deviate from U[0,1] (**Supplemental Fig. S3**), suggesting that MEMo has incorrect false positive rates. In addition, MEMo does not select “optimal” MEGS sets appropriately and typically includes many false positives (**Supplemental Fig. S4** and **Supplemental Note**). Thus, we excluded LRT-SB and MEMo from the comparison.

We simulated a mutation matrix for 54 genes in 500 samples. Among the 54 genes, mutations in 50 genes were randomly distributed. The 50 genes were classified into five groups; each group had 10 genes with mutation frequencies 1%, 5%, 10%, 20% or 30%. The simulated MEGS had 4 genes. The background mutation rates for these 4 genes were set as 1%. We simulated two types of MEGS (Fig. 3A). One had balanced mutation frequencies, i.e. all 4 genes in MEGS were mutated with the same frequency. The other had imbalanced mutation frequencies with ratio 3:1:1:1. Comparison was based on *de novo* analyses. The maximum size of MEGS was set as 8. The simulation results for MEGS with 3 genes are reported in **Supplemental Fig. S5.**

**Figure 3.**
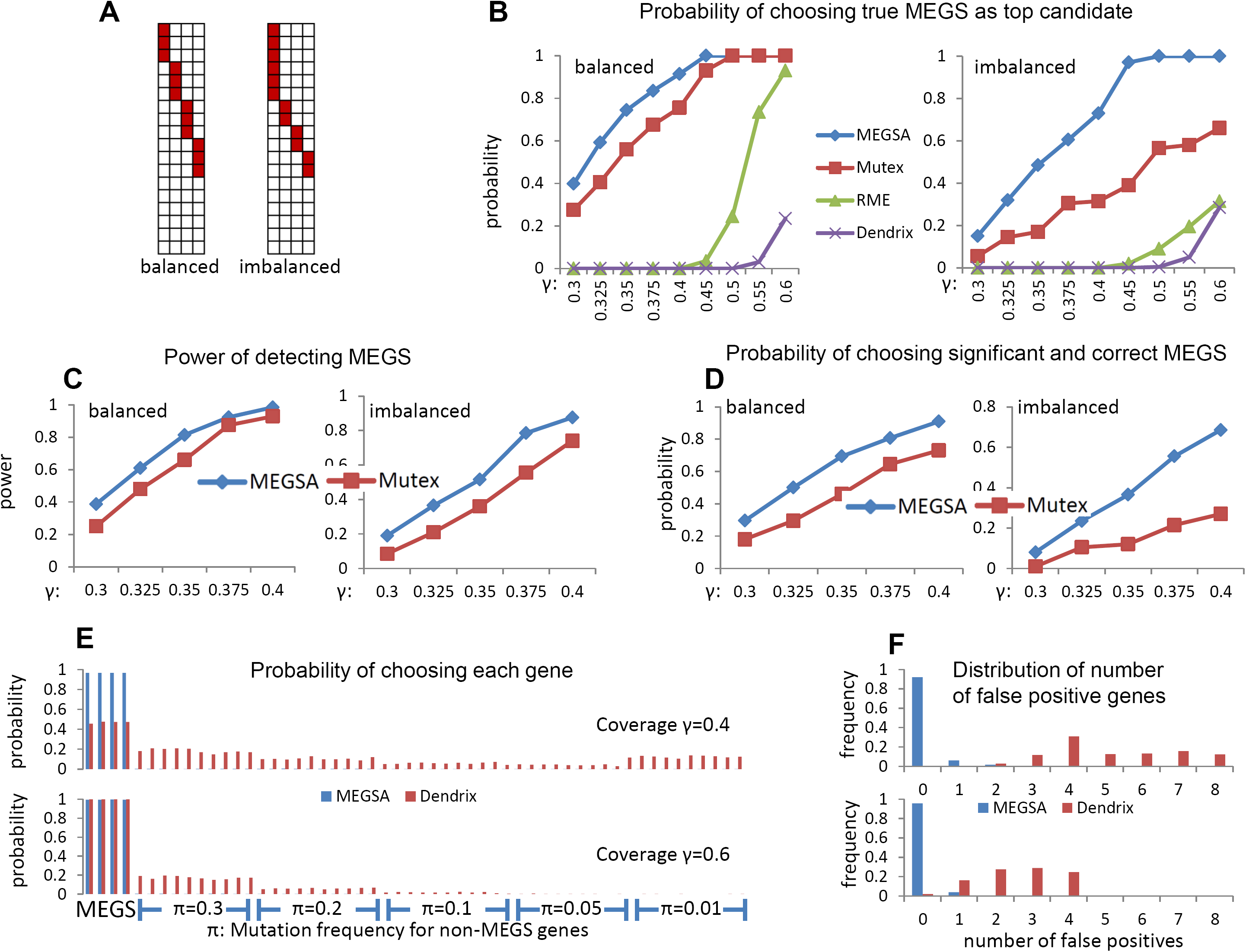
Performance comparison of methods for detecting mutually exclusive gene sets on simulated datasets. In all simulations, we have 54 genes with 50 being randomly simulated with specific mutation frequencies and 4 genes as MEGS. (A) Balanced and imbalanced MEGS with four genes. In imbalanced MEGS, the mutational frequencies have a ratio 3:1:1:1. (B) Probability of ranking the exact MEGS as the top candidate. The X-coordinate is the coverage (γ) of simulated MEGS. (C) Power of detecting MEGS using MEGSA and Mutex. (D) Probability that the identified top MEGS is statistically significant and identical to the true MEGS. Coverage (γ) of MEGS ranges from 0.3 to 0.4. (E) Probability of choosing each gene in the identified top MEGS by MEGSA and Dendrix. The top figure is based on an MEGS with coverage γ=0.4 and the bottom figure based on coverage γ=0.6. π is the mutation frequencies for the 50 non-MEGS genes. The first four are MEGS genes and the rest are non-MEGS genes. (F) The distribution of the number of falsely detected genes for the top MEGS identified by in MEGSA and Dendrix. MEGSA had few false positive genes while Dendrix detected many false positive genes.

We first compared the performance of these methods as a “scoring” method without considering the statistical significance. Thus, we calculated the probability of choosing the true MEGS identified as the top candidate for each method. Simulation results show that MEGSA performs the best for all simulations and greatly improves existing methods particularly for imbalanced MEGS (Fig. 3B). Of note, the performances are heavily impacted by the coverage of the MEGS for all methods. Dendrix has the worst performance and cannot identify the true MEGS even when the coverage is high. RME performs poorly for low coverage MEGS but reasonably well when coverage increases to 60% for balanced MEGS. Mutex outperforms RME and Dendrix.

Among Dendrix, RME, Mutex and MEGSA, only Mutex and MEGSA performed permutations to accurately evaluate overall significance (either family-wise error rate or FDR). Thus, we compared the performance of these two methods for statistically significant findings. For MEGSA, a significant finding was identified if its multiple testing corrected *P*-value <0.05. For Mutex, a significant finding was identified if FDR <0.05. A simulation was considered successful if the detected top MEGS involved any pair of the 4 genes in the simulated MEGS. The power is calculated as the proportion of “successful” simulations (Fig. 3C). A much more rigorous criterion required that the top MEGS was statistically significant and identical to the simulated MEGS (Fig. 3D). We also calculated the average number of correctly identified genes (out of 4) and number of falsely identified genes **(Supplemental Fig. S6)**. MEGSA outperforms Mutex in all comparisons. Importantly, the performance of MEGSA is superior to that of Mutex for imbalanced MEGS, which are much more frequent than balanced MEGS in real data.

Although the three methods, RME, Mutex and MEGSA, have different performances, the probability of choosing the exact MEGS increases to one when sample sizes increase to infinity, an important statistical property called “consistency”. However, the widely used Dendrix algorithm does not have this property and tends to include many false positive genes (See **Supplemental Note** for explanation). Here, we report more detailed simulation results for Dendrix, investigating the false positives in the selected top candidate. Fig. 3E reports the probability of choosing each gene based on 1000 simulations assuming coverage γ=40% (top) and γ=60% (bottom). Fig. 3F reports the distribution of the number of selected false positive genes. For example, when coverage γ=40%, in about 30% simulations, Dendrix’s top candidate includes four false positive genes. For low coverage MEGS withγ =40%, Dendrix chooses too many false positives, mostly in highly mutated genes (frequency π=30%) and lowly mutated genes (frequency π=1%). When coverage increases to 60%, Dendrix identified almost all genes in MEGS but still included many false positive genes. These simulation results suggest that, a high coverage MEGS identified by Dendrix may include multiple false positive genes. Thus, MEGS identified by Dendrix may need to be interpreted with caution. Encouragingly, MEGSA has consistently high sensitivity and low false positive rates.

Finally, we investigated the power performance of MEGSA when input genes can be partitioned into *L* modules of equal sizes by incorporating pathway information. MEGSA was applied separately to each module to generate a module-wise *P*-value. A module was statistically significant if its *P*-value < 0.05/L based on the Bonferroni correction. Under the assumption that the true MEGS is completely contained in one of the modules, the power of detecting MEGS can be substantially improved compared to *de novo* analysis that simultaneously analyzes all genes (**Supplemental Fig. S7).**

### 2.3 Analysis of The Cancer Genome Atlas (TCGA) mutation data

We analyzed non-synonymous point somatic mutations identified by whole exome sequencing for 14 cancers in TCGA with data downloaded from the data portal (https://tcga-data.nci.nih.gov/tcga/). For cancer types included in the TumorPortal website (http://cancergenome.broadinstitute.org), we included candidate driver genes reported by the website^14^ using MutsigCV^1^. Brain low grade glioma (LGG) is not reported in the TumorPortal website. Thus we identified candidate driver genes using MutsigCV^1^ and included these genes into analysis. Sample sizes, numbers of selected genes and mutational frequencies are summarized in **Supplemental Tables S2** and **S3.** For each cancer type, we performed *de novo* analysis followed by the secondary analysis incorporating KEGG pathways. For *de novo* analysis, gene sets were considered statistically significant if P< 0.05 after multiple testing based on 10,000 permutations. For KEGG-guided analysis, we derived a module-wise *P*-value for each module and declared significance by controlling FDR< 0.05. Note that MEGS from pathway-guided analyses were discarded if they were a subset of any MEGS identified in *de novo* analyses.

*De novo* analyses identified non-pairwise MEGS for acute myeloid leukemia (LAML), LGG, breast invasive carcinoma (BRCA), skin cutaneous melanoma (SKCM), head and neck squamous cell carcinoma (HNSC) and uterine corpus endometrial carcinoma (UCEC). For other cancer types, *de novo* analysis only identified pair wise MEGS. Here, we report detailed results for BRCA and LAML. The complete results are summarized in **Supplemental Table S4.**

#### 2.3.1 Analysis results for BRCA

*De novo* analyses identified 10 significant but overlapping MEGS for BRCA with 989 patients. These MEGS involved 11 genes with *TP53* involved in all MEGS (Fig. 4A). We identified five MEGS with P<10^-4^ (Fig. 4B). These MEGS were not reported by the BRCA article^15^ using MEMo^6^ that relies on functional data, emphasizing the necessity of *de novo* search.

**Figure 4.**
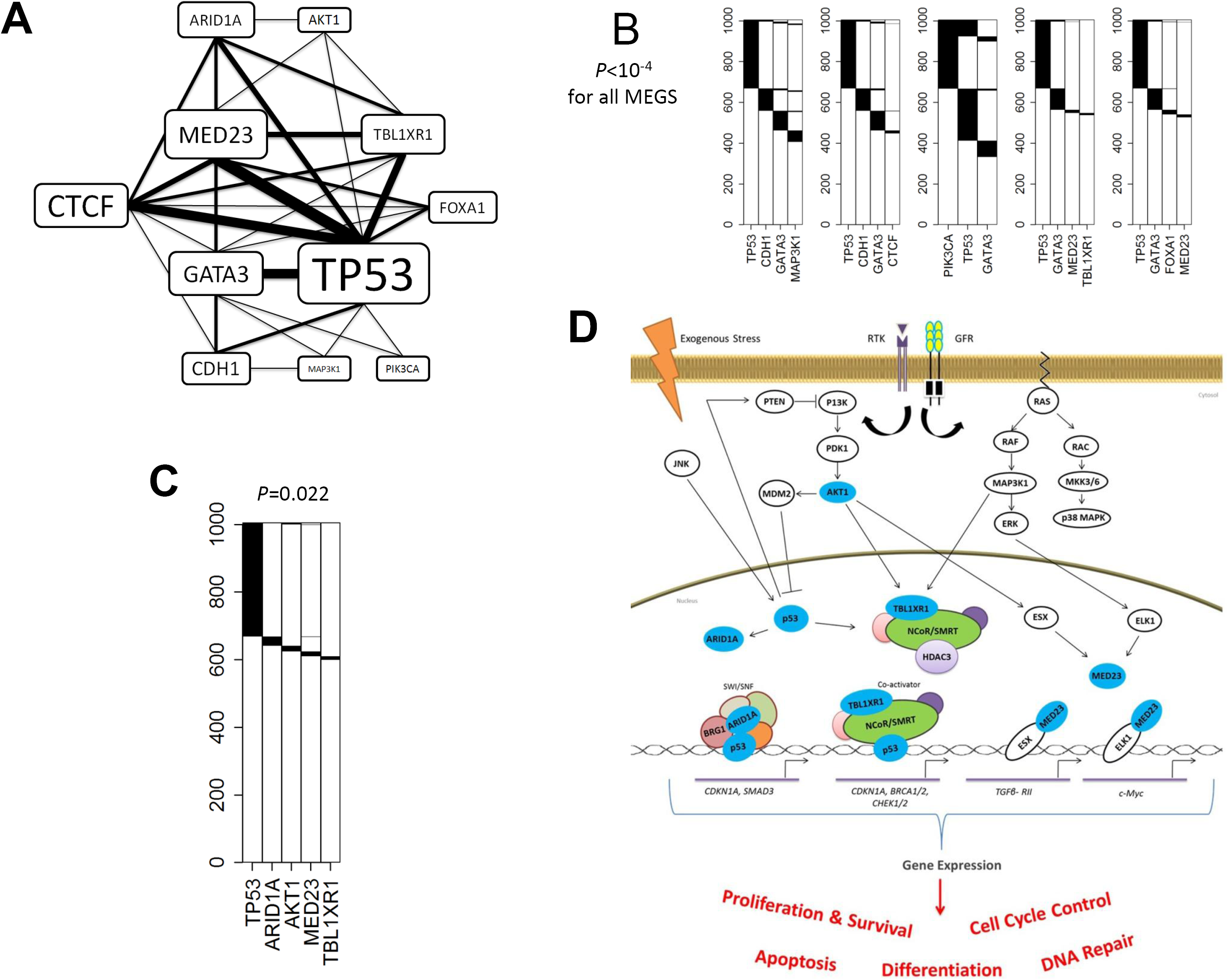
Analysis results for TCGA breast cancer whole exome sequencing data. *P*-values were adjusted for multiple testing for all reported MEGS. (A) A network constructed based on the 10 significant MEGS. Thickness of the edges and sizes of the gene labels are proportional to the times in the detected MEGS. (B) Five significant MEGS with P<10^-4^. (C) A significant MEGS with five genes. (D) Illustration showing MEGS pattern including protein products (colored blue) of *AKT1, TP53, ARID1A, MED23* and *TBL1XR1* in their relevant biological pathways. Connections including activation and interaction as well as effects on gene expression and biological processes are indicated. Components in the NCoR/SMRT and SWI/SNF complexes and the potential interaction of MED23 with p53 via the overall mediator complex are not illustrated. Connections including activation (lines with arrow) and inhibition (bar-headed lines) as well as end biological effects between the gene products are illustrated. Abbreviations: RTK, receptor tyrosine kinases; GFR, growth factor receptor.

The most significant MEGS has four genes (*TP53, CDH1, GATA3*and *MAP3K1*) and covers 59.6% of patients. E-cadherin, encoded by *CDH1,* is important in epithelial-mesenchymal transition (EMT). Morever, GATA3, p53 and MAP3K1 are related to the expression of *CDH1.* Loss of p53 represses E-cadherin expression *in vitro* as a result of *CDH1* promoter methylation^16^; GATA3 expression is correlated with E-cadherin levels in breast cancer cells^17^; and E-cadherin expression can be repressed by Snail/Slug following activation by the MAPK/ERK pathway^18^.

The largest MEGS has five genes *TP53, AKT1, ARID1A, MED23* and *TBL1XR1* (*P*=0.022) covering 40.4% of patients (Fig.s 4C and 4D). Of note, this MEGS is extremely imbalanced: all genes except *TP53* are infrequently mutated with frequency 1-2% and could not be identified by other methods, consistent with the results of simulations. TBL1XR1 belongs to and regulates the core transcription repressor complexes NCoR/SMRT^19^, and p53 gene targets may be regulated by the SMRT *in vitro* in response to DNA damage^20^. *ARID1A* encodes BAF250a, a component of the SWI/SNF chromatin-remodeling complex that directly interacts with p53^21-24^. Therefore, loss of ARID1A may have a similar effect as p53 deficiency. The mutual exclusivity between *MED23* and other genes have not been reported previously. MED23 is a subunit of the mediator complex, a key regulator of gene expression and is required for Sp1 and ELK1-dependent transcriptional activation in response to activated Ras signaling^25-28^. MED1 and MED17 directly interact with p53 ^28^, suggesting a possible connection between p53 and MED23 via the mediator complex. Also, MED23 interacts directly with the transcription factor ESX/ELF3^28^, which is downstream of AKT1 in the PI3K pathway. ESX-dependent transcription following activation by AKT is key for cell proliferation and survival. In summary, these genes have key roles in chromatin remodeling (*TBL1XR1* and *ARID1A*), gene expression regulation (*MED23* and *TP53*), and signaling (*AKT1*), and likely regulate common set of gene targets downstream of the p53, PI3K and MAPK/ERK signaling pathways that are important for cell cycle control, survival and proliferation.

Importantly, these infrequently mutated genes are unlikely to achieve high statistical significance using frequency-based driver gene test, e.g. MutSigCV^1^. In fact, in the BRCA article^15^, *MED23* and *ARID1A* were not reported as significantly mutated while *FOXA1* and *CTCF* were reported only as “near significance”. Because MutSigCV is highly sensitive to the choice of “Bagle” gene set for estimating the silent mutation rate, a very large sample size is required to replicate these findings. Given that *TP53* is a well-established driver gene, the observed mutual exclusivity provides strong and independent evidence for establishing these genes’ role as drivers.

Pathway-guided analysis identified 8 MEGS that were not detected by *de novo* analyses. Interestingly, we found that *CBFB* was mutually exclusive of *ARID1A, MED23* and *TP53.* As described above, p53 can interact with ARID1A in the SWI/SNF chromatin remodeling complex via BRG1 (see Fig. 4D for SWI/SNF complex). The transcriptional coactivator CBFB is known to interact with the tumor suppressor RUNX1, the predominant RUNX family member in breast epithelial cells^29^. RUNX1 interacts with SWI/SNF via BRG1^30^, and act as transcriptional coactivator for p53 in response to DNA damage^31^. Thus, we propose that the loss of either one of these genes would be sufficient to lead to abnormal SWI/SNF complexes and dysregulation of chromatin-related epigenetics and gene expression leading to inhibition of apoptosis.

Breast cancer is highly heterogeneous with only two genes (*TP53* and *PIK3CA*) with mutation frequencies greater than 15% **(Supplemental Table S3).** The majority of genes have mutation frequencies around 1-2%, making it difficult to identify MEGS. We successfully identified 10 significant MEGS based on *de novo* analyses and additional 8 guided by KEGG pathways. Other biological databases, e.g. functional data and Human Reference Network in MEMo^6^ or the common downstream target database in Mutex^11^, could be used in the future to guide the search of MEGS.

#### 2.3.2 Analysis results for LAML

Compared with other cancer types, AML genomes have the lowest somatic mutation rates^1^, with only 13 mutations in coding regions in average. Such a low overall (and also background) mutation rate suggests a good statistical power even with a small sample size according to our simulations **(Supplemental Fig. S2).** In fact, *de novo* analyses identified 5 distinct but overlapping significant MEGS. These significant MEGS involve 9 genes with *TP53* and *FLT3* shared by 4 MEGS (Fig. 5A). The pathway-guided search did not detect additional MEGS.

**Figure 5.**
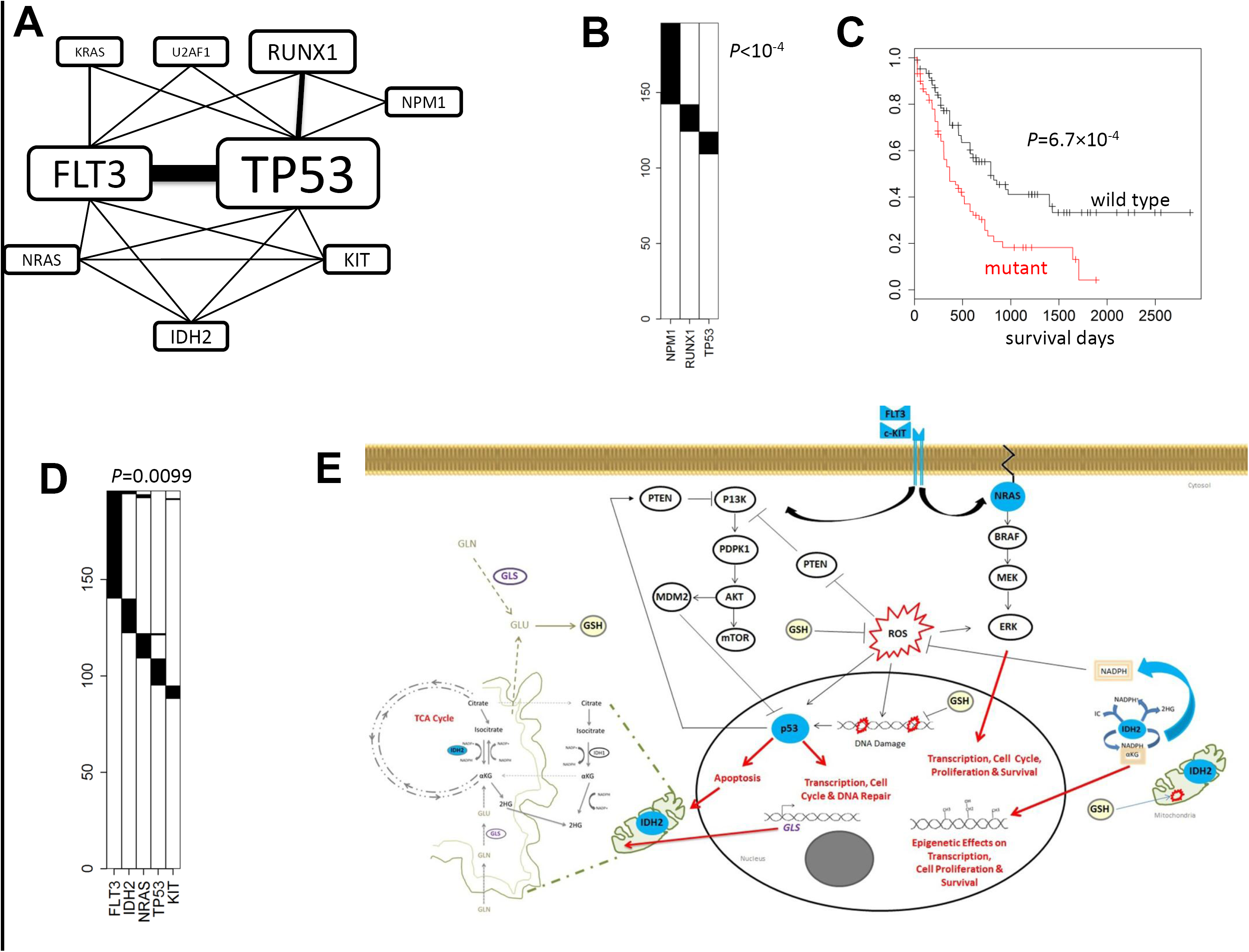
Analysis results for TCGA acute myeloid leukemia whole exome sequencing data. *P*-values were adjusted for multiple testing for all reported MEGS. (A) A network constructed based on the 5 significant MEGS. Thickness of the edges and sizes of the gene labels are proportional to the times in the detected MEGS. (B) The most significant MEGS with three genes *NPM1, RUNX1* and *TP53.* (C) The mutation status of the triplet (*NPM1, RUNX1* and *TP53*) is strongly associated with survival. (D) A significant MEGS with five genes. (E) Illustration showing MEGS pattern including protein products (colored blue) of *FLT3, IDH2, KIT, NRAS*and *TP53* in their relevant biological pathways. Connections including activation (lines with arrow) and inhibition (bar-headed lines) as well as end biological effects between the gene products are indicated. IDH2 which locates to the mitochondria is shown outside (or in-part) of the illustrated organelle for clarity and only the relevant components of glutamine (GLN) and glutathione (GSH) metabolism and TCA cycle are indicated. PI3K pathway (receptor tyrosine kinases (RTK), FLIT3 and KIT), MAPK/ERK pathway (NRAS). Abbreviations: ROS, reactive oxygen species, αKG, alpha-ketoglutarate; 2HG, 2-hydroxyglutarate; TCA, tricarboxylic acid cycle; GLN, glutamine; GLU, glutamate; and GLS, glutaminase 2; GSH (glutathione).

The most significant MEGS (Fig. 5B) has three genes *NPM1, RUNX1* and *TP53* (*P*<10^-4^), which is a subset of the top MEGS (four genes and four fusions) reported by the TCGA LAML article^32^. We further tested the association of the mutations in these three genes and their combinations with survival, adjusting for age, stage and gender. Strikingly, the strongest association was detected for the MEGS (*P*=6.7×10^-4^, Fig. 5C) but not any subset (*P*_TP53_=0.002, *P*_NPM1_=0.13, *P*_RUNX1_=0.24, P_TP53/NPM1_=0.0034, P_TP53/RUNX1_= 0.0032 and P_RUNX1_/_NPM1_=0.042), suggesting the usefulness of the MEGS for predicting clinical outcomes. Note that in the LAML article, the top MEGS included *CEBPA,* which had three (out of 13) mutations co-occurring with the triplet. In fact, including *CEBPA* into the triplet lowered the LRT statistic from 25.1 to 22.7. Thus, our model selection procedure excluded *CEBPA.* Moreover, including *CEBPA* did not significantly improve the prediction of survival (*P*=5.9×10^-4^ with *CEBPA* v.s. *P*=6.7×10^-4^ without *CEBPA*). These results suggest that the mutual exclusivity between *CEBPA* and other genes is at least suspicious and requires independent replication.

The largest MEGS (Fig. 5D) has five genes *FLT3, IDH2, KIT, NRAS*and *TP53* (*P*=0.0099), covering 55.1% of patients. This MEGS was not reported by the LAML article^32^. Fig. 5E describes important connections between function/pathways for the five gene products suggesting biological plausibility. *FLT3* and *KIT* encode receptor tyrosine kinases upstream of the PI3K and MAPK/ERK signaling pathways, and NRAS is also part of MAPK/ERK. Mutations activating theses pathways or inactivating *TP53* are common mechanisms that cancer cells use to proliferate and escape apoptosis^33^.

Interestingly, we discovered that *IDH2* belongs to this MEGS. *IDH2* encodes a mitochondrial enzyme, that converts isocitrate to α-ketoglutarate (αKG) in the tricarboxylic acid cycle and in this process produces the antioxidant nicotinamide adenine dinucleotide phosphate (NADPH), which is necessary to combat oxidative damage/stress^34,35^. Mutant IDH2 is predicted to result in: depletion of α-KG; a decrease in NADPH; production of 2-hydroxyglutarate (2-HG); and may elevate cytosolic reactive oxygen species (ROS)^36^. Mutant IDH2 can result in epigenetic effects on gene transcription (including DNA hypermethylation and histone demethylation); while loss of p53 function can result in increased expression of DNA methyltransferase 1 (DNMT1)^16,37^. Thus, a reasonable explanation for the observed mutual exclusivity between *TP53* and *IDH2* is that the loss of either protein activity may result in similar aberrant gene methylation patterns across the genome and dysregulated gene expression. We also suggest a further novel hypothesis. Depleted α-KG levels in *IDH2* mutant cells may be replenished by the conversion of glutamate to α-KG in the mitochondria^38^. The provision of glutamate in the mitochrondria is regulated by p53 via expression of the enzyme glutaminase (GLS), which also regulates antioxidant defense function in cells by increasing reduced glutathione (GSH) levels^39^. Thus, *IDH2* and *TP53* mutations are mutually exclusive because loss of both genes (or gene activity) would not be conducive to tumorgenesis or survival as a result of further depletion of αKG levels in the mitochondria and DNA damage caused by high levels of ROS. The mutual exclusivity between *IDH2* mutation and *FLT3, KIT* and *NRAS* is also biologically plausible. Mutant *IDH2* may also be linked to the activation of RAS/ERK and the PI3K pathways via ROSs, which can act as potent mitogens when apoptosis is inhibited^40^. Elevated ROS levels can activate ERKs, JNKs, or p38 and reversibly inactivated PTEN^41,42^. Thus *IDH2* mutation may be sufficient to exclusively deregulate cell proliferation and survival processes important for AML development.

## 3. Discussion

We developed a powerful and flexible framework, MEGSA, for identifying mutually exclusive gene sets (MEGS). MEGSA outperforms existing methods for *de novo* analyses and greatly improves the capability of recovering the exact MEGS, particularly for highly imbalanced MEGS. The key components of MEGSA are a likelihood ratio test and a model selection procedure. Because likelihood ratio test is asymptotically most powerful, MEGSA is expected to be nearly optimal for *de novo* search. Our algorithms can be easily adapted to other methods that integrate with external information, e.g. MEMo and Mutex, to improve performance. As an important contribution, we carefully examined the performance of existing methods. We concluded that many methods had incorrect false positive rates and poor performance for selecting optimal MEGS. Importantly, mutual exclusivity analysis may help identify infrequently mutated driver genes, as we demonstrated in the TCGA BRCA data.

MEGSA can be further improved in several ways. First, MEGSA does not consider the extremely variable somatic mutation rates across patients. Including patients with very high mutation rate may increase the background mutation rate and thus decrease the statistical power. We are currently extending MEGSA by modeling patient-specific background mutation rates. Second, MEGSA uses a multiple-path search algorithm for computational consideration and may miss findings. The Markov Chain Monte Carlo (MCMC) or the genetic algorithm may address the issue.

In the current manuscript, we analyzed TCGA non-synonymous point mutations for the purpose of testing the MEGSA algorithm. We plan to extend the analysis to include somatic copy number aberrations (SCNAs), recurrent gene fusions^32^ and epigenetic alternations. Moreover, it would be extremely interesting to restrict analysis to clonal point mutations carried by all cancer cells. Clonal mutations happen before the most recent common ancestor and are located early in the evolution tree of the tumor^43^; thus clonal mutations are likely relevant for tumorigenesis. Focusing the analysis on clonal mutations, although technically challenging^44-46^, can substantially reduce the background mutation rates and consequently improve statistical power. More importantly, this refined analysis may better reveal oncogenic pathways related with tumorigenesis.

## 4. Methods

### 4.1 A likelihood ratio statistic for testing mutual exclusivity

Suppose that a MEGS has *m* genes with mutation matrix denoted as *A*_*0*_. We assume that the *m* genes are completely mutually exclusive. A MEGS is characterized by two parameters: the coverage γ, defined as the proportion of samples covered by the MEGS, and the relative mutation frequencies *P =* (*p*_*1*_…*p*_*m*_). Background mutations are mutually independent and also independent of the MEGS mutations. We allow different background mutation rates. ∏ = (*π*_1_…*π*_*m*_) for different genes. See Fig. 2B.

For patient *i*, let (*a*_*i*1_, …, *a*_*im*_) be the observed binary mutation vector for *m* genes. Let *C*_*i*_ be a discrete binary variable. If the patient is not covered by the MEGS, *C*_*t*_ = 0. If the patient has a mutation in gene *k* in the MEGS, then *C*_*t*_ = *k*. The likelihood of observing (*a*_*i*1…._*a*_*im*_) is given by

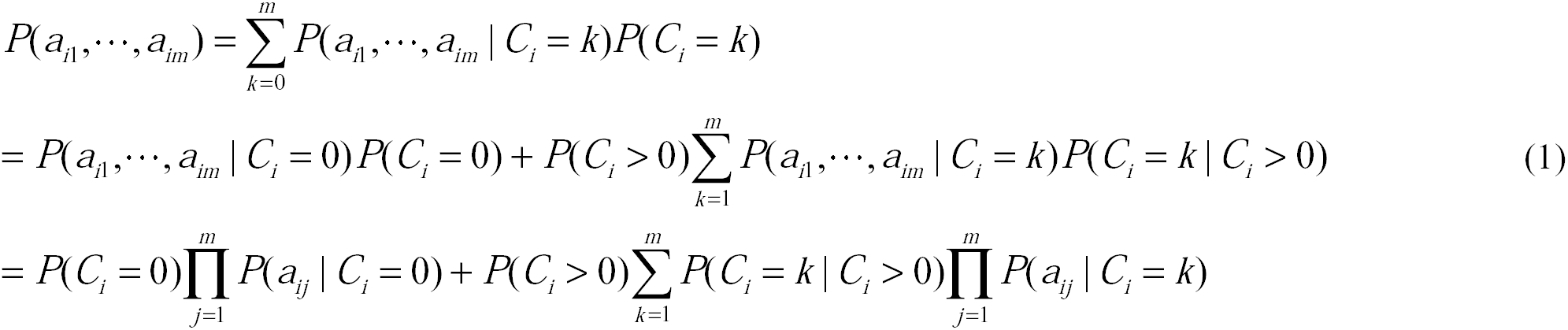

The last equation holds because mutations are independent across genes. By the definition of coverage,

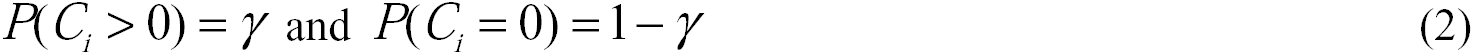

Also,

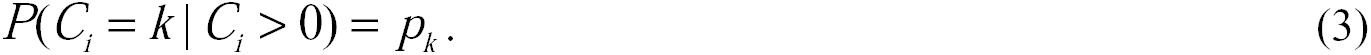

If the patient is not covered by the MEGS,

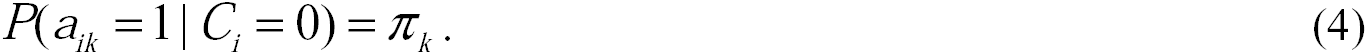

Furthermore,

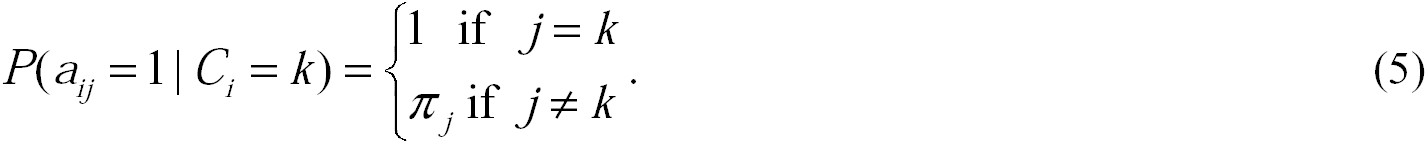

Combining (1)-(5), we have

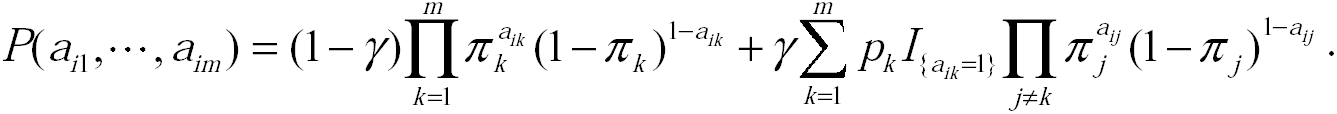

The total likelihood across *N* patients is

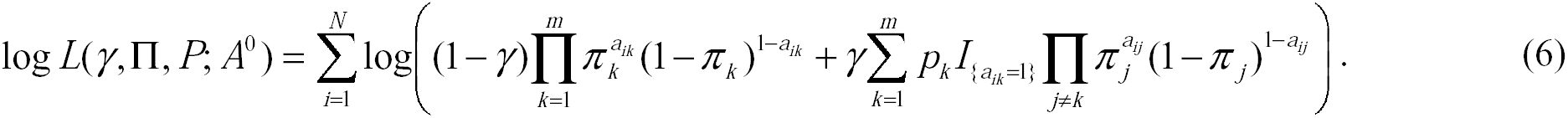

We test *H*_*0*_: **γ***=* 0 v.s. *H*_1_: **γ**> 0. *P* = (*p*_1_…. *p*_*m*_) and **∏** = (*π*_1_…*π*_*m*_) are nuisance parameters. While both parameters can be estimated under *H*_1_, *P* = (*p*_*1*_,…,*p*_*m*_) is not involved in the likelihood under *H*_0_, which causes problems in deriving asymptotic null distribution for the likelihood ratio test (LRT). To overcome this problem, we further assume that the MEGS mutation frequencies are proportional to the background mutation frequencies, i.e *p*_*k*_ ∝*π*_*k*_ Under this assumption, (6) reduces to

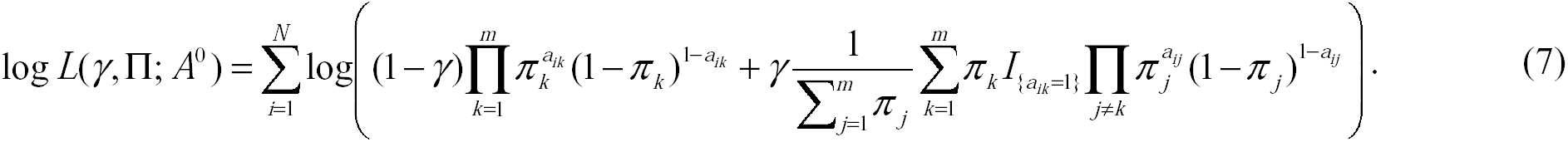

Let 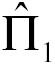 and 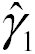, be the estimate under *H*_*1*_ and 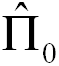 be the estimate under *H*_0_. The LRT is calculated as 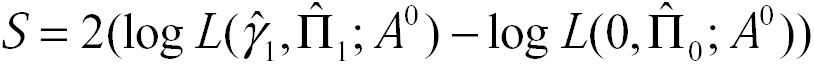 Asymptotically, LRT has a null distribution 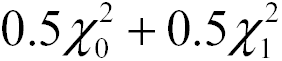 a mixture distribution with 0.5 probability at point mass zero and 0.5 probability as 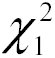.

We have two comments. First, the assumption *p*_*k*_ ∞ *γ*_*k*_ does not affect the null distribution of LRT because *p*_*k*_ is not involved in the data generation process under *H*_*0*_. However, violation of this assumption may cause power loss, which warrants further investigation. Second, the LRT in the LRT-SB method was derived based on a different data generative model, which incorrectly and unnecessarily assumed that background mutations could happen only for patients covered by the MEGS. Under this model, their likelihood function degraded as the coverage *γ* → 0, preventing them from using the standard statistical theory to derive the null distribution. To overcome this problem, they used Vuong’s^47^ method (but incorrectly) to derive an incorrect asymptotic null distribution. More details are **in Supplemental Note.**

### 4.2 Testing the global null hypothesis

Our algorithm for testing the global null hypothesis (GNH) has the following steps. (1) For *k* (*k*≤*K*), we search all gene sets of size *k* from *M* genes to test for ME using LRT and denote the minimum *P*-value as *P*_*k*_. (2) We run *T*permutations, calculate the minimum LRT *P*-value *P*_*k*_(*t*) for permutation *t*and estimate the significance (denoted as *Q_k_*) of the observed *P*_*k*_ as the proportion of simulations with *P*_*k*_(*t*) smaller than the observed *P*_*k*_. Intuitively, *Q*_*k*_ measures the significance when searching only for MEGS of size *k*. (3) Because we search for MEGS of different sizes, the overall statistic for testing GNH is defined as **θ**=min (*Q_2_,…,Q_K_*), with overall significance evaluated by permutations again.

Although conceptually straightforward, it is computationally infeasible. Finding the minimum *P*-value *P*_*k*_ even for a moderate *k* (e.g. *k*=6) is computationally very challenging and not feasible for thousands of permutations. We propose a multiple-path search algorithm to address the problem. Briefly, we calculate the *P*-values for all *M(M-1)/2* pairs of genes and choose the top *L* (e.g. *L*=10) pairs to start linear search. For the *1^th^* pair (assuming *G*_*1*_ and *G*_2_), let *q*_*2*_ (*l*) be the LRT *P*-value. Next, we calculate the LRT *P*-values for *M-2* triplets (*G_1_,G_2_,G_3_*),…, (*G_1_,G_2_,G_M_*) and choose the gene (assuming *G*_3_) with the smallest *P*-value, denoted as *q*_3_(*l*). We repeat until *q*_*K*_(*l*). For each k, we approximate *P*_*k*_ bymin_1≤*L*_ *q*_*k*_(*l*), instead of exhaustive search.

### 4.3 Identify statistically significant MEGS

Remember that we use **θ**=min (*Q_2_*…,*Q*_*K*_) as the overall statistic for testing GNH. Once GNH is rejected at level α=0.05, we need to identify all combinations of genes that reach significance. First of all, based on permutations, we can identify a cut-off *1-*. In the multiple-path search algorithm described above, for each combination of *k* genes, we transform its nominal LRT *P*-value to *Q* based on permutations and declare this gene set as significant if *Q*<*θ*_1−*α*_. This procedure may identify significant but nested putative MEGS. We designed a model selection procedure described in Fig. 1D to make a choice between nested models.

## Competing interests

The authors declare that they have no competing interest.

## Authors’ contributions

XH and JS conceived the study. XH developed software and performed numerical studies. PLH contributed to the biological connections and pathway interpretation of the findings. XH, PLH and JS drafted the manuscript. All authors reviewed and commented the manuscript.

## Acknowledgements

This study utilized the high-performance computational capabilities of the Biowulf Linux cluster at the NIH, Bethesda, MD (http://biowulf.nih.gov). The authors are supported by the National Cancer Institute intramural research program.

## References

1. Lawrence, M.S. et al Mutational heterogeneity in cancer and the search for new cancer-associated genes. Nature 499, 214–8 (2013).

2. Hodis, E. et al A landscape of driver mutations in melanoma. Cell 150, 251–63 (2012).

3. Pao, W. et al KRAS mutations and primary resistance of lung adenocarcinomas to gefitinib or erlotinib. PLoS Med 2, e17 (2005).

4. Ding, L. et al Somatic mutations affect key pathways in lung adenocarcinoma. Nature 455, 1069–75 (2008).

5. Cancer Genome Atlas Research, N. Comprehensive molecular profiling of lung adenocarcinoma. Nature 511, 543–50 (2014).

6. Ciriello, G., Cerami, E., Sander, C. & Schultz, N. Mutual exclusivity analysis identifies oncogenic network modules. Genome Res 22, 398–406 (2012).

7. Vandin, F., Upfal, E., & B.J. De novo discovery of mutated driver pathways in cancer. Genome Res 22, 375–85 (2012).

8. E. & Beerenwinkel, N. Modeling mutual exclusivity of cancer mutations. PLoS Comput Biol 10, e1003503 (2014).

9. Zhao, J., Zhang, S., Wu, L.Y. & Zhang, X.S. Efficient methods for identifying mutated driver pathways in cancer. Bioinformatics 28, 2940–7 (2012).

10. Leiserson, M.D., Blokh, D., Sharan, R. & Raphael, B.J. Simultaneous identification of multiple driver pathways in cancer. PLoS Comput Biol 9, e1003054 (2013).

11. Babur, O. et al Systematic identification of cancer driving signaling pathways based on mutual exclusivity of genomic alterations. BioRxiv (2015).

12. Miller, C.A., Settle, S.H., Sulman, E.P., Aldape, K.D. & Milosavljevic, A. Discovering functional modules by identifying recurrent and mutually exclusive mutational patterns in tumors. BMC Med Genomics 4, 34 (2011).

13. Benjamini, Y. & Hochberg, Y. Controlling the false discovery rate: a practical and powerful approach to multiple testing. Journal of the Royal Statistical Society, Series B 57 (1), 289–300 (1995).

14. Lawrence, M.S. et al Discovery and saturation analysis of cancer genes across 21 tumour types. Nature 505, 495–501 (2014).

15. Cancer Genome Atlas, N. Comprehensive molecular portraits of human breast tumours. Nature 490, 61–70 (2012).

16. Peterson, E.J., Bogler, O. & Taylor, S.M. p53-mediated repression of DNA methyltransferase 1 expression by specific DNA binding. Cancer Res 63, 6579–82 (2003).

17. Yan, W., Cao, Q.J., Arenas, R.B., Bentley, B. & Shao, R. GATA3 Inhibits Breast Cancer Metastasis through the Reversal of Epithelial-Mesenchymal Transition. Journal of Biological Chemistry 285, 14042–14051 (2010).

18. Yan, W., Cao, Q.J., Arenas, R.B., Bentley, B. & Shao, R. GATA3 inhibits breast cancer metastasis through the reversal of epithelial-mesenchymal transition. J Biol Chem 285, 14042–51 (2010).

19. Mottis, A., Mouchiroud, L. & Auwerx, J. Emerging roles of the corepressors NCoR1 and SMRT in homeostasis. Genes Dev 27, 819–35 (2013).

20. Adikesavan, A.K. et al Activation of p53 transcriptional activity by SMRT: a histone deacetylase 3-independent function of a transcriptional corepressor. Mol Cell Biol 34, 1246–61 (2014).

21. Saldana-Meyer, R. et al CTCF regulates the human p53 gene through direct interaction with its natural antisense transcript, Wrap53. Genes & Development 28, 723–734 (2014).

22. Soto-Reyes, E. & Recillas-Targa, F. Epigenetic regulation of the human p53 gene promoter by the CTCF transcription factor in transformed cell lines. Oncogene 29, 2217–2227 (2010).

23. Guan, B., Wang, T.L. & Shih, I.M. ARID1A, a Factor That Promotes Formation of SWI/SNF-Mediated Chromatin Remodeling, Is a Tumor Suppressor in Gynecologic Cancers. Cancer Research 71, 6718–6727 (2011).

24. Samartzis, E.P. et al Loss of ARID1A expression sensitizes cancer cells to PI3K- and AKT-inhibition. Oncotarget 5, 5295–303 (2014).

25. Wang, G. et al Mediator requirement for both recruitment and postrecruitment steps in transcription initiation. Molecular Cell 17, 683–694 (2005).

26. Stevens, J.L. et al Transcription control by E1A and MAP kinase pathway via Sur2 mediator subunit. Science 296, 755–758 (2002).

27. Yang, X. et al Selective requirement for Mediator MED23 in Ras-active lung cancer. Proceedings of the National Academy of Sciences of the United States of America 109, E2813–E2822 (2012).

28. Poss, Z.C., Ebmeier, C.C. & Taatjes, D.J. The Mediator complex and transcription regulation. Critical Reviews in Biochemistry and Molecular Biology 48, 575–608 (2013).

29. Chimge, N.O. & Frenkel, B. The RUNX family in breast cancer: relationships with estrogen signaling. Oncogene 32, 2121–30 (2013).

30. Decristofaro, M.F. et al Characterization of SWI/SNF protein expression in human breast cancer cell lines and other malignancies. J Cell Physiol 186, 136–45 (2001).

31. Ozaki, T., Nakagawara, A. & Nagase, H. RUNX Family Participates in the Regulation of p53-Dependent DNA Damage Response. Int J Genomics 2013, 271347 (2013).

32. Cancer Genome Atlas Research, N. Genomic and epigenomic landscapes of adult de novo acute myeloid leukemia. N Engl J Med 368, 2059–74 (2013).

33. Scholl, C., Gilliland, D.G. & Frohling, S. Deregulation of signaling pathways in acute myeloid leukemia. Semin Oncol 35, 336–45 (2008).

34. Cairns, R.A., Harris, I.S. & Mak, T.W. Regulation of cancer cell metabolism. Nat Rev Cancer 11, 85–95 (2011).

35. Cohen, A.L., Holmen, S.L. & Colman, H. IDH1 and IDH2 mutations in gliomas. Curr Neurol Neurosci Rep 13, 345 (2013).

36. Smolkova, K. & Jezek, P. The Role of Mitochondrial NADPH-Dependent Isocitrate Dehydrogenase in Cancer Cells. Int J Cell Biol 2012, 273947 (2012).

37. Lin, R.K. et al Dysregulation of p53/Sp1 control leads to DNA methyltransferase-1 overexpression in lung cancer. Cancer Res 70, 5807–17 (2010).

38. van Lith, S. et al Glutamate as chemotactic fuel for diffuse glioma cells: are they glutamate suckers? Biochim Biophys Acta. 1846(1), 66–74 (2014).

39. Jiang, P., Du, W. & Yang, X. p53 and regulation of tumor metabolism. J Carcinog 12, 21 (2013).

40. Wallace, D.C. Mitochondria and cancer. Nat Rev Cancer 12, 685–98 (2012).

41. Son, Y. et al Mitogen-Activated Protein Kinases and Reactive Oxygen Species: How Can ROS Activate MAPK Pathways? J Signal Transduct 2011, 792639 (2011).

42. Liou, G.Y. & Storz, P. Reactive oxygen species in cancer. Free Radic Res 44, 479–96 (2010).

43. Nik-Zainal, S. et al The life history of 21 breast cancers. Cell 149, 994–1007 (2012).

44. Carter, S.L. et al Absolute quantification of somatic DNA alterations in human cancer. Nat Biotechnol 30, 413–21 (2012).

45. Oesper, L., Mahmoody, A. & Raphael, B.J. THetA: inferring intra-tumor heterogeneity from high-throughput DNA sequencing data. Genome Biol 14, R80 (2013).

46. Landau, D.A. et al Evolution and impact of subclonal mutations in chronic lymphocytic leukemia. Cell 152, 714–26 (2013).

47. Vuong, Q.H. Likelihood Ratio Tests for Model Selection and Non-Nested Hypotheses. Econometrica 57 (2), 307–333 (1989).

